# Olfactory and Vomeronasal Receptor Feedback Employ Divergent Mechanisms of PERK Activation

**DOI:** 10.1101/239830

**Authors:** Ryan P Dalton, G Elif Karagöz, Jerome Kahiapo, Ruchira Sharma, Lisa E Bashkirova, David B. Lyons, Hiroaki Matsunami, Peter Walter

## Abstract

Mutually-exclusive chemoreceptor expression in olfactory and vomeronasal sensory neurons (OSNs and VSNs) enables odorant discrimination. This configuration involves chemoreceptor mediated activation of the endoplasmic reticulum (ER)-resident kinase PERK. PERK drives translation of the transcription factor ATF5 to preclude additional chemoreceptor expression. ATF5 translation is transient in OSNs but persistent in VSNs, suggesting chemoreceptor-specific modes of PERK activation. Herein, we showed that the ER-lumenal domain (LD) of PERK recognized vomeronasal receptor (VR)-derived peptides, suggesting direct PERK activation drives persistent ATF5 translation in VSNs. In contrast, PERK LD did not recognize olfactory receptor (OR)-derived peptides *in vitro*, and facilitating OR maturation *in vivo* prevented PERK activation, suggesting that ORs activate PERK indirectly through a failure to exit the ER. Importantly, impairing or prolonging ATF5 expression drove specific chemoreceptor repertoire biases. Together, these results demonstrate mechanistic divergence in chemoreceptor feedback and establish that differences in PERK activation promote qualitatively different gene regulatory results.

## Introduction

Rodents possess two olfactory organs: the main olfactory epithelium (MOE), which houses olfactory sensory neurons (OSNs) and the vomeronasal organ (VNO), which houses vomeronasal sensory neurons (VSNs). The MOE and VNO are both neurogenic, giving rise to new sensory neurons throughout the life of the animal^1–3^. The cell-surface sensory receptors expressed by OSNs/VSNs determine which ligands can activate them and inform their pattern of connectivity to the brain^4^. Therefore, receptor gene regulation during OSN and VSN development is considered to be central to the establishment of OSN/VSN cell fate.

Most OSNs either express olfactory receptors (ORs) or trace amine-associated receptors (TAARs). ORs detect a wide range of odors including those of innate importance as well as those whose valence is learned by association. TAARs detect innately-important odors such as those emitted by predators^5^. Each mouse OSN expresses only a single OR allele from a gene family of ~1100 intact genes, or a single TAAR allele from a family of 15 genes. VSNs express vomeronasal receptors (VRs) and detect pheromones and other semiochemicals including those from other species, which drive social and reproductive behaviors^6^. VSNs fall into three broad classes. Type I VSNs express a single type I VR (V1R) allele. Type II VSNs express a single type II VR (V2R) from families A, B, or D, as well as one or more V2R alleles from family C^7–10^. A third class of VSNs expresses genes from the formyl peptide receptor family. Together, these singular or highly restricted patterns of receptor expression underlie the discriminatory power of olfaction.

Monogenic or restricted OR/VR expression involves an initial process of ‘receptor gene choice’, followed by a receptor-elicited negative feedback signal that acts to prevent further receptor gene choice and to promote neuronal maturation^11–13^. VR choice is largely unstudied. By contrast, details are emerging on the mechanism of OR gene choice, which turns out to be extremely complex, involving nuclear aggregation of the OR genes, histone modification, the assembly of a multi-enhancer hub, and coordination between various transcriptional activators^14–18^. OR translation in the ER initiates feedback by activating the ER-resident kinase PERK, which controls one branch of the unfolded protein response (UPR)^19^, a highly-conserved homeostatic signaling pathway communicating the ER folding status to the nucleus to maintain homeostasis^3,11,12,19–22^. Upon activation, PERK drives phosphorylation of the eukaryotic initiation factor eIF2α, inhibiting global translation initiation to decrease ER protein folding burden^23,24^. In developing OSNs/VSNs, eIF2α phosphorylation also drives the selective translation of *Atf5* mRNA, a transcription factor that then promotes neuronal maturation^25^. In OSNs, ATF5 prevents further OR gene choice and stabilizes OR expression^19^.

Following VR choice in VSNs, ATF5 is translated persistently and in areas corresponding to both immature and mature VSNs^26,27^. In contrast, in OSNs ATF5 is only translated transiently, at the onset of OR expression, despite continuing presence of OR, *Perk*, and *Atf5* mRNA^19^,^26^. This difference in PERK activation dynamics and subsequent ATF5 translation suggests that PERK activation is context-dependent and that ORs and VRs may differ mechanistically in how they activate PERK.

At least two models could explain how receptors activate PERK. In the first model, receptors are direct PERK ligands. This direct binding model is supported by recent findings on the structurally-related sensor IRE1. The crystal structure of the IRE1 sensory domain displays an architecture similar to the major histocompatibility complex (MHC) peptide-binding groove^28^. IRE1’s lumenal domain directly interacts with peptides and unfolded proteins, leading to its oligomerization and activation^29,30^. While direct PERK activation by unfolded proteins has not been demonstrated, given its structural resemblance to IRE1 it is plausible that receptors activate PERK by maintaining ‘unfolded’ regions that act as PERK ligands.

In an alternative model, receptors activate PERK indirectly, through a failure to fold properly and traffic from the ER, which could result in sequestration of chaperones and could induce general protein misfolding. This indirect model is supported by experiments showing that ORs and VRs fail to traffic from the ER when expressed heterologously. Moreover, deletion of <u>r</u>eceptor <u>t</u>ransporter <u>p</u>roteins 1 and 2 <u>(RTP</u>1/2), which facilitate the ER exit and plasma membrane targeting of ORs, prolongs ATF5 translation in OSNs. Importantly, ATF5 directly binds the *Rtp1* promoter regulating RTP1 synthesis. *Rtp1/2* transcription downstream of ATF5 translation could act to terminate further ATF5 translation through either competition with PERK for OR binding sites or through facilitating OR folding and trafficking.

Here, we set out to determine (1) how ORs and VRs activate PERK, (2) why PERK activation is transient in OSNs but persistent in VSNs, and (3) how different modes of PERK activation could regulate chemoreceptor feedback programs. To this end, we tested the two models discussed above both with biochemical assays and in mouse lines engineered for this purpose. Our findings reveal the mechanistic basis of PERK activation by ORs and VRs and demonstrate that PERK activation dynamics constitute an important signal in cell fate acquisition.

## Results

### VRs display PERK interacting peptides

First we set out to determine how ORs activate the PERK branch of the UPR and why PERK activation is only transient during OSN maturation. To test whether ORs display potential sites that can be directly recognized by PERK’s ER-lumenal domain (LD), we generated peptide arrays using a series of 18-mer peptides that tile the entire length of the well-studied OR OLFR1507 (also known as MOR28) in 3 amino acid steps and probed these peptide arrays with purified PERK LD. As a control, we generated a peptide array for the V1RB2, a monogenically-expressed V1R, as well as the broadly-expressed family C V2R, VMN2R6^10^. *V1rb2* cannot be stably expressed by OSNs when used to replace an OR coding sequence, suggesting that VRs and ORs may employ somewhat different feedback mechanisms^31,32^. An N-terminal lumenal OR region bound PERK LD very weakly, while the sequences that contain hydrophobic amino acids from the neighboring transmembrane domains displayed strong PERk LD binding (**Figure 1A**). In contrast, two ER-lumenal V1RB2 regions interacted strongly with PERK LD on the peptide array, as did a number of lumenal regions derived from VMN2R6 (**Figure 1B, C**). To confirm these observations, we evaluated these peptides in solution using fluorescence anisotropy binding assays, which verified tight binding of VR (K_D_ = 6.6 ± 0.8 μM and K_D_ = 6.1 ± 1.7 μM) but not OR (K_D_ =250.3 ± 1.5 μM) peptides to PERK LD (**Figure 1D,E**). Peptide arrays do not take into account three-dimensional structure of proteins as well as the possible interactions in the context of full-length receptors, both of which might contribute to the interaction of the receptors with PERK LD. However, these results suggested an attractive model that could explain differences in PERK activation and dynamics in ORs versus VRs that we tested in our subsequent experiments.

**Figure 1.**
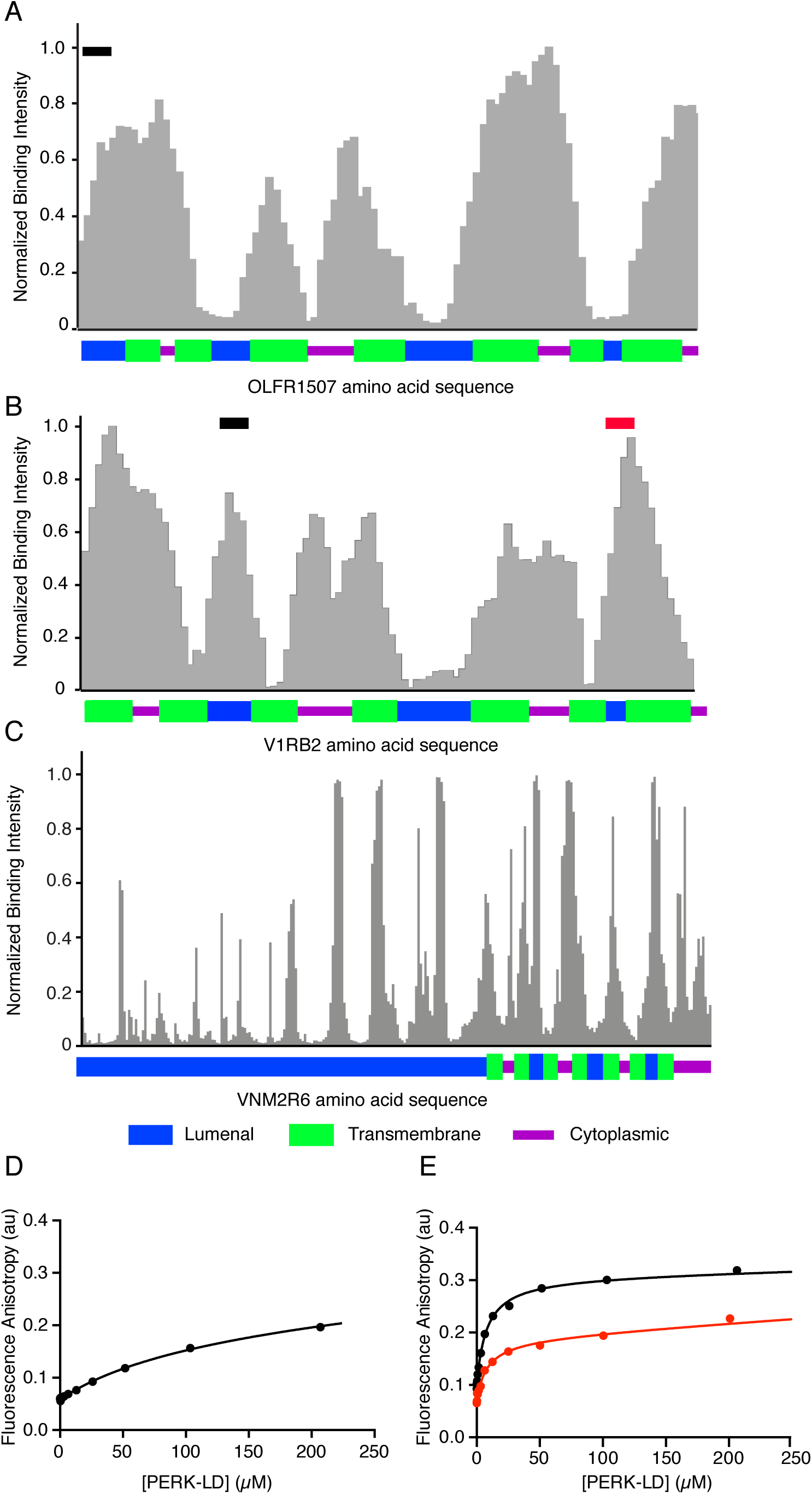
PERK’s ER-sensor domain binds distinct peptides derived from VRs. **(A)** Quantification of the peptide arrays derived from Olfactory Receptor OR1507 for binding of PERK Lumenal Domain (LD). The contribution of each amino acid from Olfactory Receptor OR1507 to the binding of PERK LD was calculated by averaging the intensity of all the spots containing that amino acid. The binding score is plotted against the amino acid sequence of OR1507. The topology of the receptor is indicated at the bottom of the graph, where blue, green, and purple bars depict lumenal, transmembrane, and cytoplasmic domains of the protein. **(B)** Quantification of the peptide arrays derived from Vomeronasal Receptor V1RB2 for binding of PERK LD. The contribution of each amino acid from Vomeronasal Receptor V1RB2 to the binding of PERK LD was calculated as in (A). The binding score is plotted against the amino acid sequence of V1RB2. The topology of V1RB2 is indicated as in (A). The black and red bars show the binding score of the peptides used in the fluorescence anisotropy experiments. **(C)** Quantification of the peptide arrays derived from Vomeronasal Receptor VMN2R6 for binding of PERK LD. The contribution of each amino acid from VMN2R6 to PERK LD binding was calculated as in (A). The topology of V1RB2 is indicated as in (A). **(D)** Fluorescence anisotropy measurements show weak binding of PERK LD to OR1507 derived peptide. The black bar shows the binding score of the peptide used in the fluorescence anisotropy experiments with K_D_ =250.3 ± 1.5 μM. **(E)** V1RB2 derived peptides V1RB2-1 (black) and V1RB2-2 (red) bind PERK LD with comparable affinity of K_D_ = 6.6 ± 0.8 μM and K_D_ = 6.1 ± 1.7 μM respectively, measured by fluorescence anisotropy experiments.

### ORs Activate PERK through a Failure to Exit the ER

As discussed above, OR expression could activate the UPR, including PERK, by overloading the ER at the onset of their translation. Alternatively, if there is an interaction between PERK and ORs in the context of full-length OR, RTP1/2 might compete with PERK for OR binding sites upon PERK-induced *Rtp1/2* expression. In both scenarios, *Rtp1/2* expression would relieve the ER burden and prevent further UPR activation. This mechanism could account for the transient nature of ATF5 translation observed in OSNs^19^. To determine whether inefficient OR transport out of the ER is responsible for transient PERK activation, we changed the timing of RTP1/2 expression. To this end, we generated a transgenic mouse line that expresses high levels of RTP1/2 under the control of a synthetic promoter (hereafter, *tetO-Rtp)*. This promoter is activated by the tetracycline transactivator protein (tTa)^33^. We have two tTa driver lines capable of driving *tetO-Rtp* expression. In the first, *tTa* is expressed under the control of the G protein *Gng8*, which is expressed coincident with OR choice (hereafter, *Gng8-tTa)* in OSNs^21,34^. In the second, *tTa* is expressed under the control of a marker of mature OSNs, *OMP* (hereafter, *OMP-tta)*^21^ VSNs also express these genes and with similar timing, allowing the use of this transgenic strategy in both the MOE and the VNO^35^.

By crossing *tetO-Rtp* mice to *Gng8-tTa*; *OMP-tTa* mice, we were able to overexpress RTP1/2 beginning at OR choice and persisting in mature OSNs. In these mice, there was very low ATF5 immunoreactivity in the MOE as compared to control animals, suggesting reduced PERK activity. We also observed reduced expression of the mature OSN (mOSN) marker ADCY3 (**Figure 2A-B**). ADCY3 was restricted to only the most apical OSNs similar to *Atf5-/-* MOEs^19^. We also observed striking changes in animals in which *tetO-Rtp* is expressed only under the control of *Gng8-tTa*. In these animals, ATF5 translation was shifted apically, similar to ADCY3 expression, which was limited to the most apical OSN layers (**Figure 2C**). OR expression has been previously shown to be required and sufficient for ATF5 translation^19^. Our transgenic *Rtp1/2* data now provide crucial context for those findings: the presence or absence of RTP1/2 in the ER appear to determine whether or not OR expression drives ATF5 translation. Importantly, these data readily explain the puzzling finding that ATF5 translation is transient. ORs are initially expressed into an ER environment mostly lacking RTP1/2. Once PERK has been activated and ATF5 has been translated, the level of RTP1/2 increases. RTP1/2 then facilitate the trafficking of the ORs, decreasing the protein folding load in the ER. Finally, given that VR trafficking is independent of RTP1/2, we predicted that the timing or degree of ATF5 translation in the VNO should not be modified in our transgenic animals. Indeed, we observed no defect in ATF5 translation in transgenic VNOs (**Figure 2D**).

**Figure 2.**
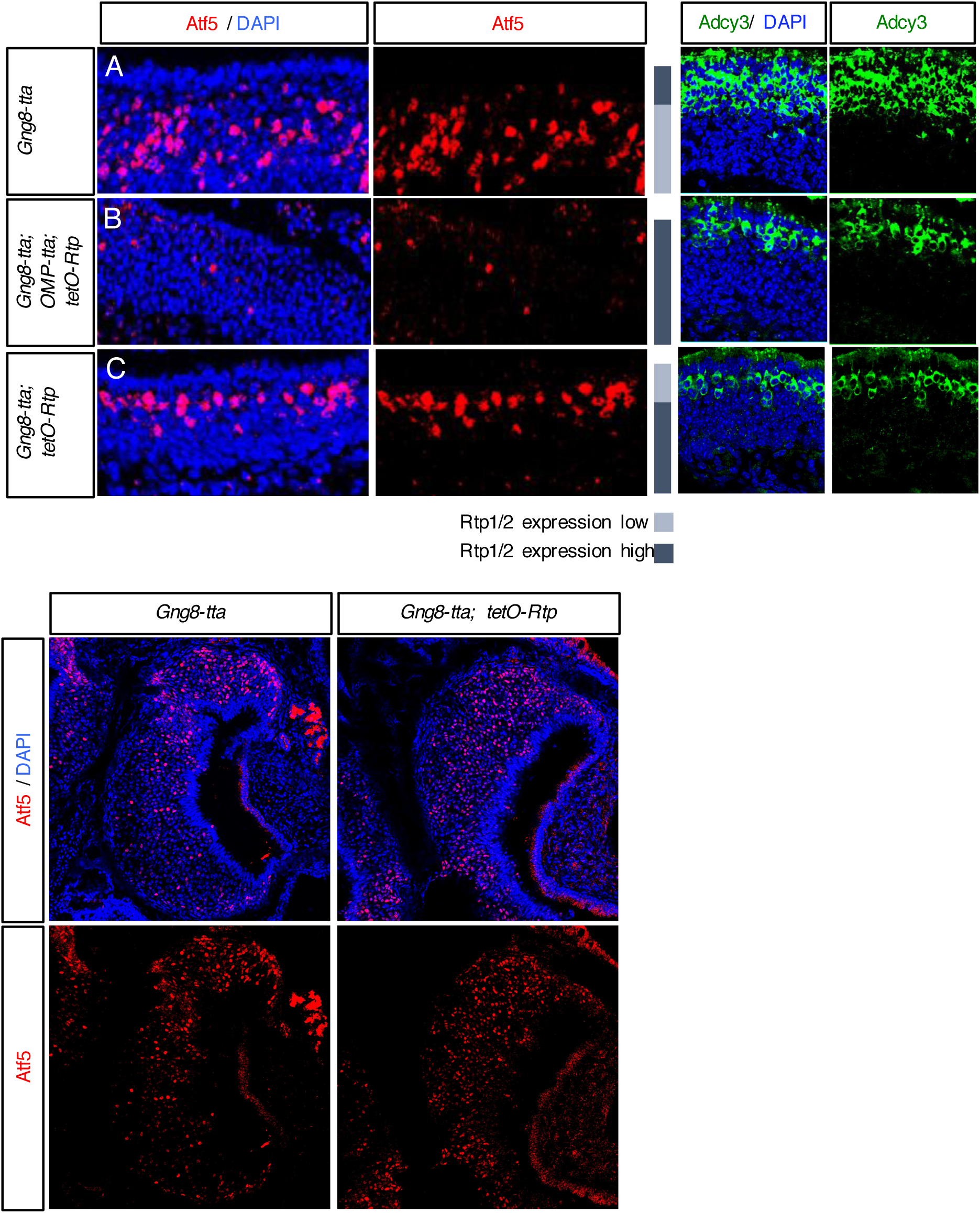
ORs activate PERK through a failure to exit the ER. **(A)** Representative coronal sections from postnatal day 14 (P14) *Gng-8tta* control animals revealed that ATF5 immunoreactivity was mainly in more basal regions of the MOE, appearing coincident with OR choice. ADCY3 protein was mutually-exclusive with ATF5 protein, labeling 4 to 5 layers of mature OSNs. **(B)** When the *tetO-Rtp1/2* transgene is placed under the control of both *Gng-8tta* and *OMP-tTa* in littermates, ATF5 protein was almost entirely lost, and ADCY3 was greatly restricted. When this transgene was only expressed under *Gng-8tta* control, coincident with OR choice, ATF5 protein was shifted apically, as was ADCY3 protein **(C)**. Inset gray bars show anticipated levels of RTP1/2 expression in control and transgenic animals. **(D)** The vomeronasal organ of an adult *Gng-8tta* control animal compared to a *Gng-8tta; tetO-Rtp1/2* animal. ATF5 immunoreactivity is broad, labeling areas both corresponding to immature and mature VSNs.

### Prolonged ATF5 Expression Results in a TAAR-type Cell Fate

The finding that the duration of ATF5 translation is controlled by the level of RTP1/2 expression suggested the interesting possibility that the mechanism, duration, and/or amplitude of PERK activation and ATF5 translation could coordinate distinct downstream transcriptional programs. For example, if OR expression were not stabilized until ATF5 translation terminated, then receptors such as VRs that persistently activate the UPR would drive gene switching in OSNs. In addition, smaller changes in ATF5 translation could coordinate expression of specific signaling molecules, chaperones, or other markers of OSN subtypes. As an initial test of this hypothesis, we looked for receptor expression biases in mouse mutants lacking *Adcy3 (Adcy3-/-)*. These animals have been shown to exhibit prolonged ATF5 translation and have increased OR gene switching, consistent with a model of OR feedback in which termination of ATF5 translation is an important signal for the stabilization of OR expression^16^. Compared to control animals, *Adcy3-/-* mice showed a striking increase in the expression of TAARs (**Figure 3A**)^36,37^. As with ORs, TAARs are expressed monogenically and monoallelically; thus if the increase in TAAR expression results from prolonged ATF5 translation, then TAAR expression should also increase in RTP1/2 double-knockout mice, which exhibit prolonged ATF5 translation^38^. Indeed, we found that these mice showed an increase in TAAR expression compared to controls (**Figure 3A**).

**Figure 3.**
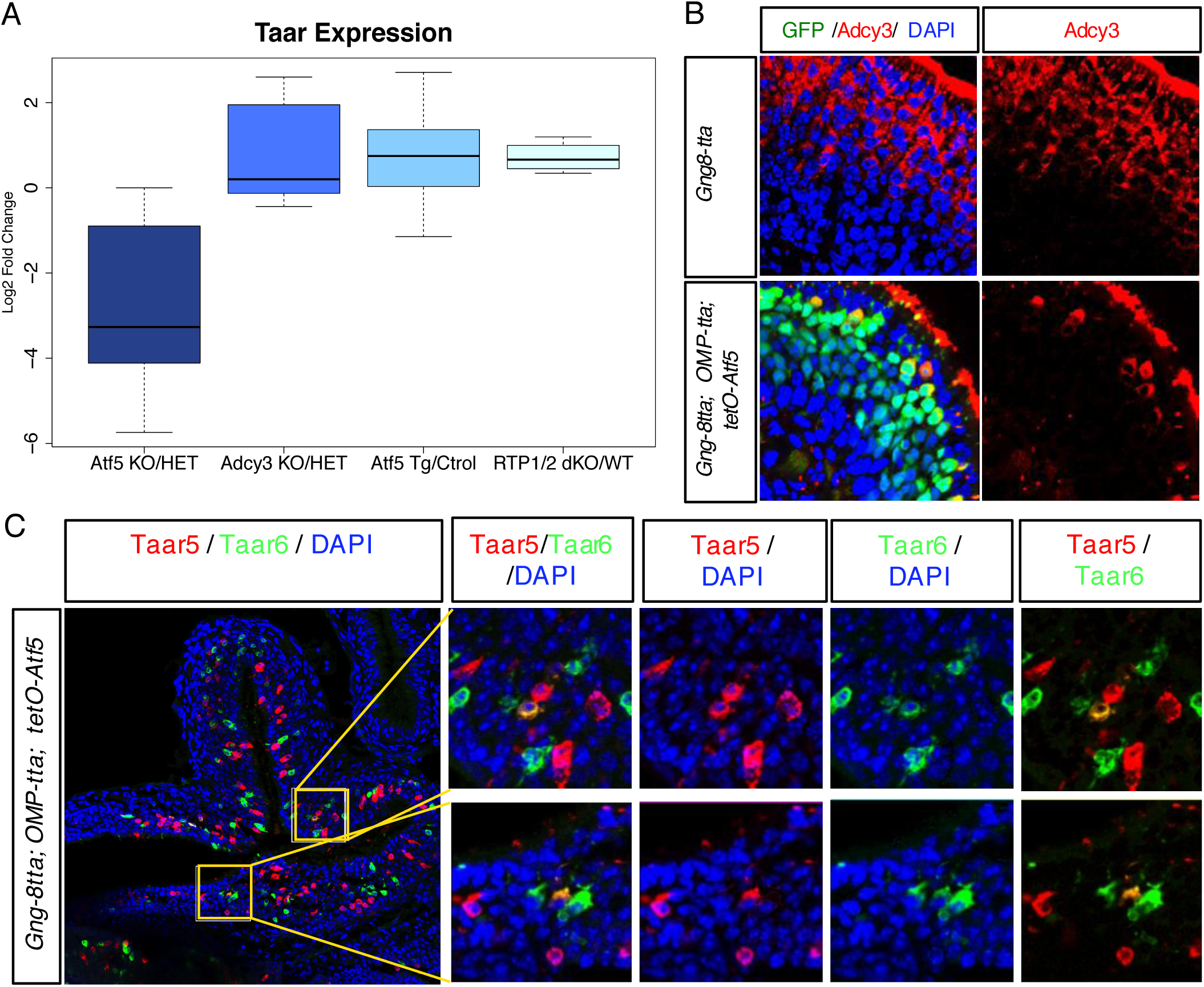
Prolonged ATF5 expression results in a TAAR-type cell fate. **(A)** Representative coronal sections showed that expression of *tetO-Atf5* under the control of *Gng8-tta* and *OMP-tta* resulted in a severe reduction in ADCY3+ cells (bottom) compared to control animals (top). Many transgene-expressing cells were ADCY3+ (bottom left panel). **(B)** Average log2 RPKM from mRNA-seq experiments for detected TAAR genes in adult *Atf5-/-* vs. *Atf5+/-, Adcy3-/-* vs. *Adcy3+/-, Gng8-tta; OMP-tta; tetO-Atf5* versus control and *Rtp1/2-/-* versus control animals. Mean log2 ratios are - 2.789, .774, .731, and .717 respectively. **(C)** Immunofluorescence for TAAR5 and TAAR6 in an adult *Gng8-tta; OMP-tta; tetO-Atf5* animal. The two insets each include an example of a cell co-expressing TAAR5 and TAAR6.

These data did not demonstrate a direct role for ATF5, as prolonged ATF5 translation in the mutant animals simply reflects prolonged activation of PERK. PERK controls translation of various other mRNAs. To test directly whether prolonged ATF5 expression increased TAAR expression, we employed the previously-published *Atf5* transgenic mouse line described above (*tetO-Atf5*)^19^. Expression of *tetO-Atf5* under the control of both *Gng8-tTa* and *OMP-tTa* prevented the expression of mature OSN markers, such as ADCY3 (**Figure 3B**). These animals also showed an increase in TAAR expression comparable to that observed in the *Adcy3* and *Rtp1/2* knockout mice (**Figure 3A**). Strikingly, we observed cells that expressed at least two TAAR genes in *tetO-Atf5* mice (**Figure 3C** and insets). The coexpression of TAAR genes in single OSNs has not been previously observed. These data indicated that prolonged ATF5 translation drives a TAAR-type cell fate in OSNs. Thus, our findings in combination with published work^16,38^ support a model in which OSN maturation and OR gene stability are not established until ATF5 translation in OSNs is terminated. In addition, the time course or amplitude of ATF5 translation appears itself to be acting as an important cue in the refinement of OSN identity, driving a TAAR-type cell fate.

### An *Atf5*-independent OSN Cell Type

As we showed that prolonged ATF5 translation drives biases in cell fate decisions, we next asked whether these biases are also observed in the absence of *Atf5*. In *Atf5-/-* mice, OSN maturation is dramatically impaired. However, a few cells differentiate successfully^19^. As has been previously reported^19^, these cells display a characteristically regular spatial orientation. This provided an initial clue that they could represent a distinct cell type that was enriched by loss of *Atf5*. To determine the identity of these cells, we sought to purify them from *Atf5-/-* mice. To accomplish this, we crossed the *Atf5-/-* animals to animals expressing green fluorescent protein (GFP) under the control of the mature OSN marker *OMP* (hereafter, *OMP-GFP)*^39^. By dissecting the MOE of these animals and using fluorescence-activated cell sorting (FACS), we isolated mature OSNs from *Atf5-/-* animals. RNA was isolated from this cell population and used for transcriptomic analysis by RNAseq. As a control, we also isolated *OMP-GFP+* cells from *Atf5+/-* animals for further RNAseq analyses. By comparing *Atf5+/-; OMP-GFP* and *Atf5-/-; OMP-GFP* sequencing results, we were able to identify specific subsets of receptors or signaling molecules that were enriched in either data set. The *Atf5-/-* data showed a decrease in canonical OR signaling molecules such as *Adcy3* and *Cnga2* compared to control. In contrast, we observed a dramatic increase in expression of the guanylate cyclase *Gucy1b2* in *Atf5-/-* compared to control. We also observed an increase in expression of the transcription factor *Emx1*, in these OSNs (**Figure 4A**). *Gucy1b2*, which is thought to be part of a non-canonical OR signaling pathway, has recently been implicated in low-oxygen sensing^40,41^, and *Emx1* was recently shown to be coexpressed with several guanylate cyclases^42^. However, it has not yet been demonstrated what receptors *Gucy1b2+* cells express or whether these receptors are distinct from those expressed by other OSNs. Our RNAseq analysis identified a small number of ORs that were highly enriched in the *Atf5-/-* MOE (**Figure 4B**). These receptors therefore potentially represent the subclass of receptors expressed by the *Gucy1b2+* cells. To confirm an increase in expression of these ORs in *Atf5-/-* animals, we performed RNA in situ hybridization, using probes against the identified receptors. As shown in **Figure 4C, t**he expression of *Olfr309*, which was among the receptors we identified, was dramatically enhanced in *Atf5-/-* animals compared to control. In summary, *Atf5* deletion offers an example of chemoreceptor expression bias that is complementary to what we observed in three models of ATF5 overexpression. It also revealed a potential molecular framework for the establishment of the *Gucy1b2+* type cell fate.

**Figure 4.**
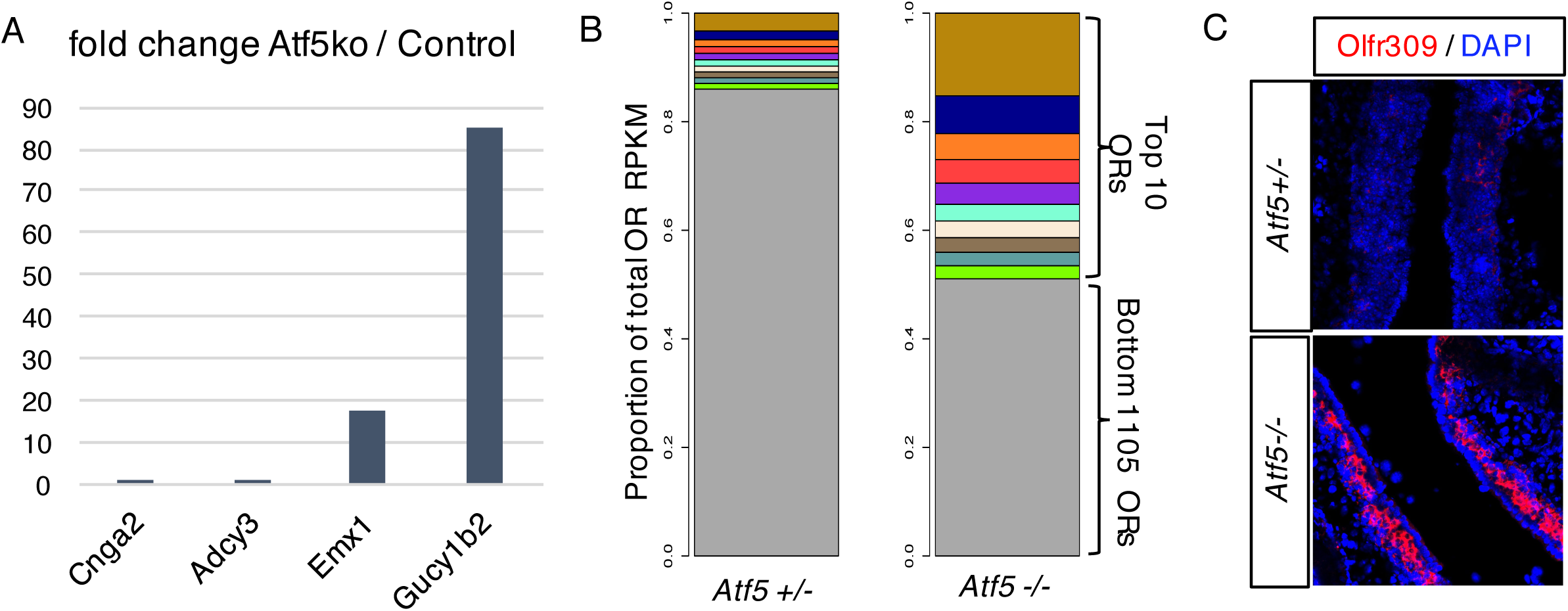
Loss of *Atf5* results in receptor bias and *Gucy1b2+* cell bias. **(A)** Fold change from mRNA-seq for *Cnga2, Adcy3, Emx1*, and *Gucy1b2* in sorted *oMPiGFP; Atf5-/-* versus sorted *OMPiGFP; Atf5+/-* animals. **(B)** OR diversity as a measure of total OR read RPKM. Top 10 most-highly expressed OR genes are labeled at top in color as proportion of RPKM, with remaining ~1100 ORs in gray. The proportion of reads in *Atf5-/-* sorted cells is nearly 50% for the top 10 ORs, as opposed to ~17% in control. **(C)** RNA *in situ hybridization* for *0lfr309*, one of the 10 most-enriched OR genes in *OMP-GFP; Atf5-/-*, in *Atf5-/-* and *Atf5+/-* adult animals.

## Discussion

With this work, we revealed the mechanistic basis for how ORs and VRs activate feedback, a fundamental feature in the development of the olfactory system. We provided insight into the puzzling observation that ORs only transiently activate PERK to induce *Atf5* translation. Our data suggest that in the case of ORs, ER folding homeostasis is restored upon OR- and ATF5-driven expression of the OR transporters RTP1/2. Once ATF5 levels have declined due to efficient trafficking of OR out of the ER, OSN maturation is completed and OR gene stability becomes finalized (**Figure 5A-C**).

**Figure 5.**
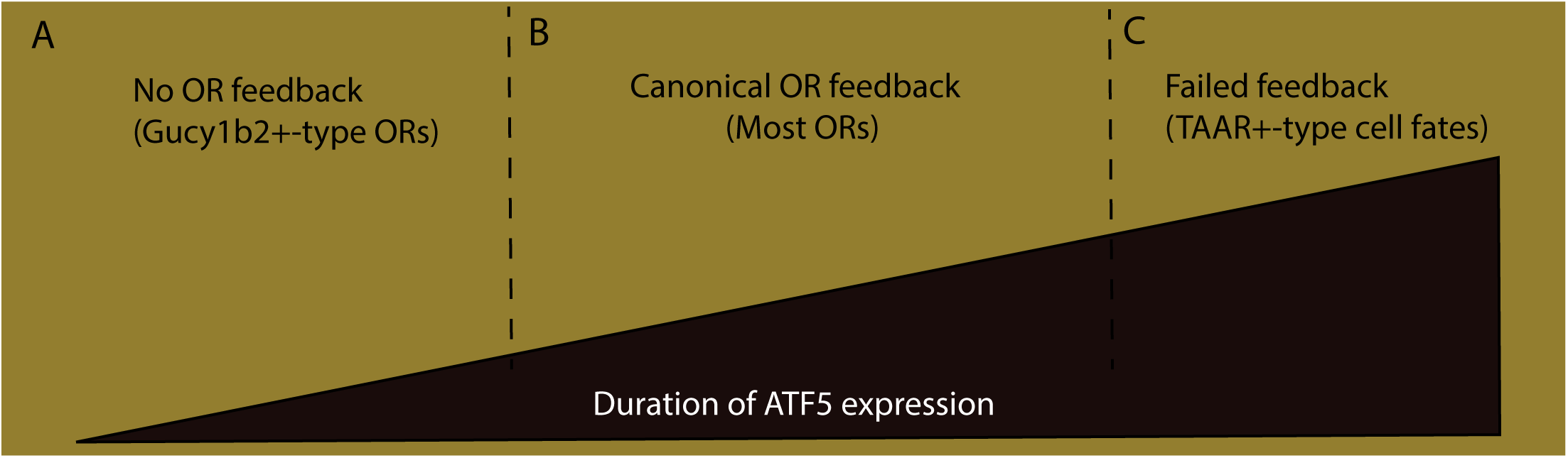
The duration of ATF5 translation influences cell fate outcomes. **(A-C)** The duration of ATF5 translation appears to influence cell fate. Cells that fail to activate ATF5 translation assume cell fates associated with *Gucy1b2* expression and a subset of ORs **(A)**. Cells that activate an intermediate level of ATF5 translation and then resolve ER homeostasis account for the majority of OSNs and constitute the ‘canonical’ OR feedback pathway-employing cells **(B)**. Prolonged ATF5 translation results in ‘failed feedback’ and TAAR expression **(C)**.

The output of the feedback pathway in OSNs appears to depend on the ratio of OR/RTP protein and the efficiency by which RTP1/2 transport ORs from the ER. In this sense, OR feedback is similar to canonical models of UPR function, as it acts to restore ER homeostasis. A key difference is that in the case of the OSN, the homeostatic adjustments appear to accommodate mainly OR protein. This should allow OSNs to discriminate between OR-driven PERK activation, which RTP1/2 would ameliorate, and general protein misfolding, which RTP1/2 would not. Coordinating OR appearance with changes to the ER folding environment has fascinating consequences. If the chosen OR is only weakly transcribed, for example if the gene fails to be recruited to a multi-enhancer hub, low levels or even absence of of RTP1/2 may sufficiently traffic the OR to prevent PERK activation. This scenario would result in gene switching, which has been observed for OR transgenes expressed at low levels. On the other hand, this could allow the beta-2 adrenoreceptor to activate feedback when expressed from an OR locus. OSNs also accommodate the opposite situation, in which an OR cannot be efficiently trafficked. This could be due to the development of an ER-retention motif, or *bona fide* misfolding, for example. In these cases, PERK activation would be prolonged, also resulting in gene switching. Thus, translation of ATF5 may act as a molecular hourglass. Only ORs that drive the correct level and duration of PERK activation will have their expression stabilized. For a chosen OR to drive ATF5 translation to the degree that OR stability results, the OR must therefore i) be expressed sufficiently to first activate UPR and ii) productively traffic with RTP1/2 to then relieve the UPR.

In extreme cases, receptor repertoires can be biased, as we observed both with *Atf5* deletion and with the three models of ATF5 overexpression described above. A detailed sequencing analysis could likely reveal the extent to which these receptor repertoires overlap, potentially uncovering additional biases. Both *Atf5* deletion and ATF5 overexpression result in gene switching, and it is therefore fascinating that we observed such different biases in receptor expression. These biases are most consistent with a model in which UPR duration aids in the molecular refinement of OSN identity. This is consistent with a recent report demonstrating a surprisingly dynamic gene regulatory response during chronic ER stress ^43^. Whether the gene expression program identified in this report helps to enact the TAAR or *Gucy1b2-type* OSN gene expression programs is a fascinating outstanding question.

By contrast to our findings in the MOE, we found that the PERK LD can recognize distinct sites in V1Rs and V2Rs, suggesting that these receptors might directly interact with PERK. The finding that ATF5 translation was prolonged in both apical and basal areas of the VNO containing mature VSNs supports this direct activation model. These findings also may explain an interesting variation between MOE and VNO feedback signals. Substitution of an OR coding sequence with a VR and *vice versa* would result in UPR induction in both types of sensory neurons. However, prolonged UPR activation would result in stable receptor expression only in VSNs, in agreement with the outcomes of these substitution experiments^20,32,44^ and in agreement with the demonstration that *Olfr692* is expressed in the VNO^45^. It therefore appears that, unlike in OSNs, termination of ATF5 translation is not required for stabilization of VR expression. How V1R and V2R regulation differ, and whether their feedback pathways help confer these differences, remains to be studied. Of particular interest is the coexpression of some V2R subclasses and *H2-Mv* genes. *H2-Mvs* form a family of nonclassical class I major histocompatibility complexes (MHC), which were suggested to contribute to VR function, yet their exact role remains unknown. Coexpression of *H2-Mv* genes with V2Rs may contribute to ligand sensitivity via enhanced V2R trafficking. Alternatively, *H2-Mvs* may perform a function more analogous to other MHCs, requiring ‘loading’ with VR peptides prior to ER export. Sequential expression of V2R subclasses and *H2-Mvs* could suggest that the *H2-Mvs* may themselves be targets of an initial feedback signal, similar to *Rtp1/2*.

Our OR and VR feedback models stem from the biochemical interrogation of only a small number of receptors. A high throughput analysis of every OR and VR, or of the binding regions identified herein, may reveal additional differences between chemoreceptor subclasses. This analysis could also identify the specific motifs, if they exist, that allow VRs to activate PERK directly and ORs to bypass such activation. However, our fundamental observation is that different chemoreceptors activate PERK differently, resulting in variable duration of ATF5 translation and, thus, distinct transcriptional consequences. In sum, our data provide mechanistic insight into OR feedback, VR feedback, and the development of various olfactory subsystems. These findings suggest that structural variation in receptors has direct consequences on the transcriptional regulation of these receptors and the molecular identity of the neurons that express them, generating additional selective constraints to the evolution of chemoreceptors. Finally, our results support a general role for PERK in the diversification of sensory neurons.

## Acknowledgments

We would like to acknowledge the members of the Walter, Lomvardas, and Matsunami laboratories for critical comments on this manuscript and Nataliya Zyma, Silke Nock, and Catrina Carey for technical support. We also wish to thank Gilad Barnea for providing antibodies raised against TAARs.

## Materials and Methods

### Mice and Strains Used

All mice were housed in standard conditions with a 12-hour light/dark cycle and access to food and water in accordance with University of California IACUC guidelines. All strains were maintained on a mixed genetic background. The following mouse lines have been previously described: *Gng8-tta and OMP-tta*^21^ *OMP-GFP*^46^, Atf5^19^, *Adcy3*^16^, *tetO-Atf5*^19^, and *Rtp1/2* knockout^38^. The *tetO-Rtp1/2* line was generated by restriction cloning. Briefly, the *Rtp1* and *Rtp2* coding sequences and an *EGFP* gene were placed downstream of a tta-inducible promoter, in tandem with two IRES elements: *tetO-Rtp1-IRES-Rtp2-IRES-EGFP*, which we refer to in shorthand as *tetO-Rtp1/2*. This line was generated by random insertion at the Gladstone transgenic gene targeting core facility at UCSF.

### Immunofluorescence and RNA *in situ* hybridization

Immunofluorescence (IF) was performed as previously described^15,16,19^. Briefly, tissue was either fixed in 4% PFA in phosphate-buffered saline (PBS) for 30 minutes and then washed in PBS for 30 minutes, sucrose protected and embedded in OCT; or was directly dissected into OCT for more sensitive antibodies. 14 μm sections were air-dried for 10 minutes, fixed in 4% PFA in PBS for 10 minutes, washed for 3 × 5 minutes in PBS + .1% Triton-X (PBST), blocked for 1 hour in 4% donkey serum in PBST, then incubated with primary antibodies under coverslips overnight at 4°C. The following day, slides were washed for 3 x 15 minutes in PBST and then incubated with Alexa dye-conjugated secondary antibodies and 4’,6-diamidino-2-phenylindole (DAPI) at concentrations of 1:1000 under cover slides. Slides were then washed for 3 × 15 minutes in PBST and mounted with Vectashield for imaging. Imaging was performed on Leica 700-series laser scanning confocal microscopes. The following antibodies were used: goat anti-Atf5 (SCBT SC-46934, dilution 1:250); rabbit anti-ADCY3 (SCBT SC-588, dilution 1:300); anti-TAAR5 and anti-TAAR6 (from Gilad Barnea, dilution 1:1000). RNA *in situ* hybridization was performed as previously described^16^.

### mRNA-Seq

RNA for RNA-seq libraries was isolated from either whole MOE as described previously^14^ or from FAC-sorted cells. Sequencing libraries were prepared in-house using standard methods and Nugen Ovation RNA-seq reagents. Single-end 50bp reads were sequenced on Illumina Hiseq v4 at the Genomic Services lab of Hudson Alpha in New York City, NY. *Gng8-tta; OMP-tta tetO-Atf5* and *Gng8-tta; OMP-tta* control libraries were sequenced in duplicate, with, respectively, 44M, 21M, 16M, and 30M reads. Reads were normalized using Cuffnorm and all analysis was performed in R.

### Reagents

Synthetic signal peptides were ordered from Gen Script at > 95% purity. Each peptide had a 5’-fluorescein isothiocyanate (FAM) tag. Soluble peptides used in this study are, OR-1: 5-FAM-MEKAVLINQTSVMSFR, V1rb2-1: 5-FAM-MFMPWGRWNSTTCQSLIYLHR and V1rb2-2: 5-FAM-LKFKDCSVFYFVHIIMSHSYA

### Protein Purification

To express mouse PERK GST-PERK (aa 33-417) or 6x-His-PERK (aa 97-417), PERK was cloned into a pGEX4T1 vector to create a fusion protein containing N-terminal GST or to pet28a vector with N-terminal 6 × His fusion. The plasmids were transformed into *Escherichia coli* strain BL21DE3 codon plus RIPL cells (Agilent Technologies). **E. coli** BL21-DE3* RIPL cells expressing the constructs were grown in Luria-Bertani broth (LB) at 37°C until OD 600 = 0.6 and expression of proteins was then induced with 0.3 mM isopropyl β-D-1 thiogalactopyranoside (IPTG) at 21°C overnight. Cells were harvested and resuspended in lysis buffer (50 mM HEPES pH 7.2, 400 mM NaCl, 4 mM 1,4-Dithiothreitol (DTT) (or 5 mM β-mercaptoethanol if His-TRAP HQ column was used, (GE Healthcare)), and Roche protease inhibitor cocktail). Resuspended cells were lysed with the Avestin EmulsiFlex-C3 at 16,000 psi. After lysis, the supernatant was collected following centrifugation for 40 minutes at 30,000 × g. Supernatant was batch bound to glutathione sepharose resin (GE Healthcare) for 2 hours. The columns were washed with 20 column volumes of lysis buffer (50 mM HEPES pH 7.2, 400 mM NaCl, 4 mM DTT, Roche protease inhibitor cocktail) and eluted with 3 column volumes of elution buffer (50 mM HEPES pH 7.2, 150 mM NaCl, 4 mM DTT+ 20 mM Glutathione). The 6x-His-PERK constructs were purified on His-TRAP HQ column (GE, Healthcare), washed with 25 column volumes of wash buffer (50 mM HEPES pH 7.2, 400 mM NaCl, 20 mM imidazole, 5 mM β-mercaptoethanol, Roche protease inhibitor cocktail) and eluted with a gradient of 20 mM to 500 mM imidazole in wash buffer. Eluates of both affinity purifications were diluted with buffer at 50 mM HEPES pH 7.2 to 50 mM NaCl and applied to a MonoQ ion exchange column and eluted with a linear gradient from 50 mM to 1 M NaCl. The protein was then further purified on a Superdex 200 10/300 gel filtration column equilibrated with buffer A ( 25 mM HEPES pH 7.2, 150 mM NaCl, 4 mM DTT). The concentration of protein was determined using the predicted extinction coefficient at 280 nm.

### Peptide arrays

Peptide arrays were purchased from the MIT Biopolymers Laboratory. The tiling arrays were composed of 18mer peptides tiled along the sequence of OLFR1507, V1RB2, and VMN2R6 proteins with a 3 amino acid shift at a time. The arrays were incubated in methanol for 10 minutes, then in binding buffer (50 mM HEPES pH 7.2, 150 mM NaCl, 0.02% Tween-20, 2 mM DTT) three times for 10 minutes. After washing, the arrays were incubated for 1 hour at room temperature with 500 nM GST-PERK LD. The arrays were washed again for three times for 10 min in binding buffer to remove the unbound protein. Using a semi-dry transfer apparatus, the bound protein was transferred to a nitrocellulose membrane and detected with Abcam α-GST S tag antibody. The contribution of each amino acid to PERK LD binding was calculated as described previously (Gardner and Walter, 2011).

### Fluorescence Anisotropy

Binding affinity of PERK LD to 5’-FAM-labeled peptides was measured by the change in fluorescence anisotropy on a Spectramax-M5 plate reader with λex = 485 nm and λem = 525 nm with increasing concentrations of PERK LD. 50-100 nM of fluorescently labeled peptide was used in each reaction. The reaction volume of each data point was 20 μL. The measurements were done in 384-well, black flat-bottomed plates after incubation of peptide with protein for 30 min at 25^o^ C. The curves were fit using Graphpad Prism.

**Figure 1-Figure supplement 1.**
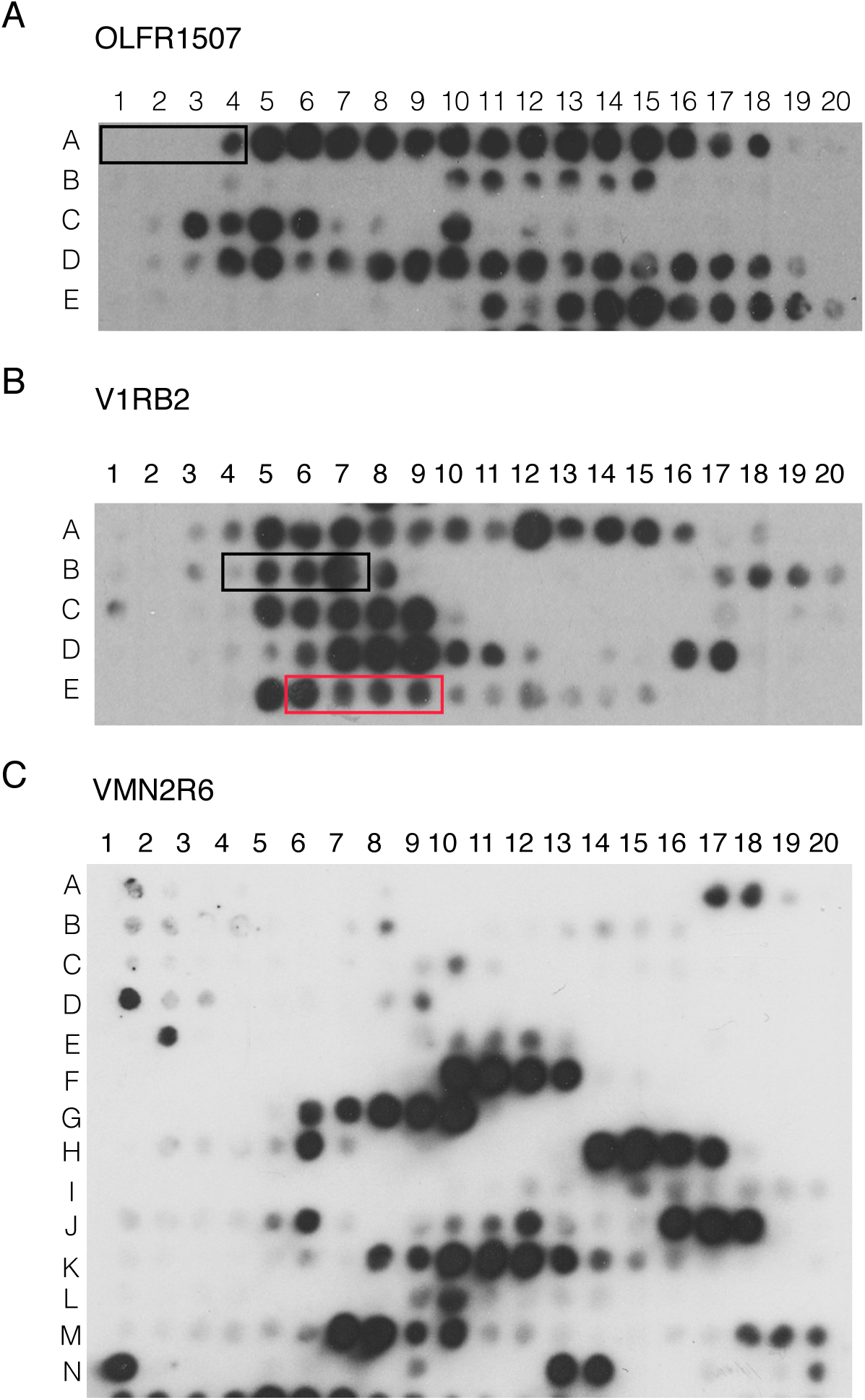
PERK luminal domain binds select peptides derived from VRs and ORs. The peptide arrays tiled with 18mer peptides derived from Olfactory Receptor OR1507 (A) and Vomeronasal Receptors V1RB2 (B) and VMN2R6 (C) were probed with purified PERK lumenal domain. The red and black boxes depict the peptide sequences that are used in the fluorescence anisotropy assays in Fig. 1.

## References

1. Graziadei, P. P. C. Cell dynamics in the olfactory mucosa. Tissue and Cell 5, 113–131 (1973).

2. Graziadei, P. P. C. & Monti Graziadei, G. A. Neurogenesis and neuron regeneration in the olfactory system of mammals. III. Deafferentation and reinnervation of the olfactory bulb following section of thefila olfactoria in rat. Journal of Neurocytology 9, 145–162 (1980).

3. Dalton, R. P. & Lomvardas, S. Chemosensory receptor specificity and regulation. Annu. Rev. Neurosci. 38, 331–349 (2015).

4. Mombaerts, P. et al. Visualizing an olfactory sensory map. Cell 87, 675–686 (1996).

5. Ferrero, D. M. et al. Detection and avoidance of a carnivore odor by prey. Proceedings of the National Academy of Sciences 108, 11235–11240 (2011).

6. Dulac, C. & Axel, R. A novel family of genes encoding putative pheromone receptors in mammals. Cell 83, 195–206 (1995).

7. Ishii, T. & Mombaerts, P. Coordinated coexpression of two vomeronasal receptor V2R genes per neuron in the mouse. Molecular and Cellular Neuroscience 46, 397–408 (2011).

8. Berghard, A. & Buck, L. B. Sensory transduction in vomeronasal neurons: evidence for G alpha o, G alpha i2, and adenylyl cyclase II as major components of a pheromone signaling cascade. Journal of Neuroscience 16, 909–918 (1996).

9. Ryba, N. J. & Tirindelli, R. A new multigene family of putative pheromone receptors. Neuron 19, 371–379 (1997).

10. Silvotti, L., Cavalca, E., Gatti, R., Percudani, R. & Tirindelli, R. A recent class of chemosensory neurons developed in mouse and rat. PLoS ONE 6, e24462 (2011).

11. Shykind, B. M. et al. Gene switching and the stability of odorant receptor gene choice. Cell 117, 801–815 (2004).

12. Lewcock, J. W. & Reed, R. R. A feedback mechanism regulates monoallelic odorant receptor expression. … Academy of Sciences of the United … (2004).

13. Serizawa, S. et al. Negative Feedback Regulation Ensures the One Receptor-One Olfactory Neuron Rule in Mouse. Science 302, 2088–2094 (2003).

14. Magklara, A. et al. An epigenetic signature for monoallelic olfactory receptor expression. Cell 145, 555–570 (2011).

15. Clowney, E. J. et al. Nuclear aggregation of olfactory receptor genes governs their monogenic expression. Cell 151, 724–737 (2012).

16. Lyons, D. B. et al. An epigenetic trap stabilizes singular olfactory receptor expression. Cell 154, 325–336 (2013).

17. Markenscoff-Papadimitriou, E. et al. Enhancer interaction networks as a means for singular olfactory receptor expression. Cell 159, 543–557 (2014).

18. Monahan, K. et al. Cooperative interactions enable singular olfactory receptor expression in mouse olfactory neurons. Elife 6, 1083 (2017).

19. Dalton, R. P., Lyons, D. B. & Lomvardas, S. Co-opting the unfolded protein response to elicit olfactory receptor feedback. Cell 155, 321–332 (2013).

20. Capello, L., Roppolo, D., Jungo, V. P., Feinstein, P. & Rodriguez, I. A common gene exclusion mechanism used by two chemosensory systems. Eur. J. Neurosci. 29, 671–678 (2009).

21. Nguyen, M. Q., Zhou, Z., Marks, C. A., Ryba, N. J. P. & Belluscio, L. Prominent Roles for Odorant Receptor Coding Sequences in Allelic Exclusion. Cell 131, 1009–1017 (2007).

22. Serizawa, S. Negative Feedback Regulation Ensures the One Neuron-One Receptor Rule in the Mouse Olfactory System. Chemical Senses 30, i99–i100 (2005).

23. Harding, H. P., Zhang, Y. & Ron, D. Protein translation and folding are coupled by an endoplasmic-reticulum-resident kinase. Nature 397, 271–274 (1999).

24. Walter, P. & Ron, D. The unfolded protein response: from stress pathway to homeostatic regulation. Science 334, 1081–1086 (2011).

25. Wang, S.-Z., Ou, J., Zhu, L. J. & Green, M. R. Transcription factor ATF5 is required for terminal differentiation and survival of olfactory sensory neurons. Proc. Natl. Acad. Sci. U.S.A. 109, 18589–18594 (2012).

26. Dalton, R. P. Shared Genetic Requirements for Atf5 Translation in the Vomeronasal Organ and Main Olfactory Epithelium. (2017). doi:10.1101/224980

27. Nakano, H. et al. Activating transcription factor 5 (ATF5) is essential for the maturation and survival of mouse basal vomeronasal sensory neurons. Cell Tissue Res. 363, 621–633 (2016).

28. Credle, J. J., Finer-Moore, J. S., Papa, F. R., Stroud, R. M. & Walter, P. On the mechanism of sensing unfolded protein in the endoplasmic reticulum. Proceedings of the National Academy of Sciences 102, 18773–18784 (2005).

29. Gardner, B. M. & Walter, P. Unfolded proteins are Ire1-activating ligands that directly induce the unfolded protein response. Science 333, 1891–1894 (2011).

30. Karagöz, G. E. et al. An unfolded protein-induced conformational switch activates mammalian IRE1. Elife 6, 53 (2017).

31. Rodriguez, I., Feinstein, P. & Mombaerts, P. Variable patterns of axonal projections of sensory neurons in the mouse vomeronasal system. Cell 97, 199–208 (1999).

32. Feinstein, P. & Mombaerts, P. A contextual model for axonal sorting into glomeruli in the mouse olfactory system. Cell 117, 817–831 (2004).

33. Gossen, M. & Bujard, H. Tight control of gene expression in mammalian cells by tetracycline-responsive promoters. Proceedings of the National Academy of Sciences 89, 5547–5551 (1992).

34. Ryba, N. J. & Tirindelli, R. A novel GTP-binding protein gamma-subunit, G gamma 8, is expressed during neurogenesis in the olfactory and vomeronasal neuroepithelia. J. Biol. Chem. 270, 6757–6767 (1995).

35. Tirindelli, R. & Ryba, N. J. The G-protein gamma-subunit G gamma 8 is expressed in the developing axons of olfactory and vomeronasal neurons. Eur. J. Neurosci. 8, 2388–2398 (1996).

36. Liberles, S. D. & Buck, L. B. A second class of chemosensory receptors in the olfactory epithelium. Nature 442, 645–650 (2006).

37. Johnson, M. A. et al. Neurons expressing trace amine-associated receptors project to discrete glomeruli and constitute an olfactory subsystem. Proc. Natl. Acad. Sci. U.S.A. 109, 13410–13415 (2012).

38. Sharma, R. et al. Olfactory receptor accessory proteins play crucial roles in receptor function and gene choice. Elife 6, 1083 (2017).

39. Potter, S. M. et al. Structure and emergence of specific olfactory glomeruli in the mouse. J. Neurosci. 21, 9713–9723 (2001).

40. Bleymehl, K. et al. A Sensor for Low Environmental Oxygen in the Mouse Main Olfactory Epithelium. Neuron 92, 1196–1203 (2016).

41. Omura, M. & Mombaerts, P. Trpc2-expressing sensory neurons in the mouse main olfactory epithelium of type B express the soluble guanylate cyclase Gucy1b2. Mol. Cell. Neurosci. 65, 114–124 (2015).

42. Parrilla, M., Chang, I., Degl'Innocenti, A. & Omura, M. Expression of homeobox genes in the mouse olfactory epithelium. Journal of Comparative Neurology 524, Spc1–Spc1 (2016).

43. Guan, B.-J. et al. A Unique ISR Program Determines Cellular Responses to Chronic Stress. Molecular Cell 68, 885–900.e6 (2017).

44. Wang, F., Nemes, A., Mendelsohn, M. & Axel, R. Odorant receptors govern the formation of a precise topographic map. Cell 93, 47–60 (1998).

45. Nakahara, T. S. et al. Detection of pup odors by non-canonical adult vomeronasal neurons expressing an odorant receptor gene is influenced by sex and parenting status. BMC Biol. 14, 12 (2016).

46. Li, J., Ishii, T., Feinstein, P. & Mombaerts, P. Odorant receptor gene choice is reset by nuclear transfer from mouse olfactory sensory neurons. Nature 428, 393–399 (2004).

